# Cardiac activity impacts spinal cord excitability. A Call to Return to the Roots

**DOI:** 10.1101/2025.10.15.682433

**Authors:** Nikolay Syrov, Polina Morozova, Anna Popova, Elizaveta Melashenko, Renat Takhirov, Alfia Mustafina, Asma Benachour, Lev Yakovlev, Mikhail Knyshenko, Alexander Kaplan

## Abstract

The heart continuously shapes neural processing and behavior through cardiac-brain interactions. While cortical excitability fluctuations and their role in cardiac-dependent cognitive and sensorimotor phenomena have been extensively studied, the temporal dynamics and contribution of spinal excitability oscillations across the cardiac cycle remain poorly characterized. In this study, we examine whether motor evoked potentials elicited by magnetic stimulation of the spinal cord are modulated by the cardiac cycle phase in healthy participants. Real-time adapting ECG-triggered stimulation enabled precise targeting of five phases across the cardiac cycle. Spinal excitability was significantly phase-dependent, with MEPs peaking during late diastole. MEPs amplitude was also found to be modulated by preceding cardiac intervals, where shorter intervals predict stronger diastolic facilitation. These findings establish that spinal excitability is rhythmically modulated by the cardiac cycle, potentially through blood pressure-mediated mechanisms. Notably, the diastolic facilitation observed here contrasts with previously reported motor cortex excitability profiles, indicating a non-synchronous cardiac modulation of spinal versus cortical excitability. These results may benefit neuromodulation approaches for motor and psychiatric disorder treatment and emphasize the critical importance of including spinal measures in future heart-brain interaction studies.

## Introduction

The heart constantly transmits information to the brain, where it is processed and influences both neuronal bodily functioning (Engelen et al., 2024; Khoshnoud et al., 2024; Mussini et al., 2025). Although nervous system monitoring of cardiac activity occurs predominantly unconsciously, cognitive and sensorimotor performance fluctuates with the cardiac cycle (Al et al., 2020; Engelen et al., 2024; Zaccaro et al., 2025). While several candidate mechanisms may explain heart-driven fluctuations in brain function—including baroreceptor effects through the solitary nucleus tract and reticular formation, direct cardiac afferents, tactile peripheral receptors, and vascular pulsatility effects on mechanosensitive channels—the specific pathways responsible for these effects have not been definitively established (De Falco et al., 2024; Engelen et al., 2024; Jammal Salameh et al., 2024; Larra et al., 2020). Transcranial magnetic stimulation (TMS) provides noninvasive assessment of cortical excitability through electromyography (EMG) motor evoked potentials (MEPs) and TMS-evoked potentials (TEPs), offering insights into brain-heart interaction mechanisms. Al et al. (2023) demonstrated that M1 cortical excitability differs across the cardiac cycle phases, with higher excitability during the systole which decreases linearly through the diastole phase (Al et al., 2023). Additionally, (Paci et al., 2024) found stronger GABA-mediated intracortical inhibition during systole compared to diastole phase, supporting enhanced motor control during this phase (Galvez-Pol et al., 2022; Konttinen et al., 2003; Palser et al., 2021). At the same time, numerous other TMS studies have reported no robust cardiac cycle effects on M1-MEP amplitudes (Bianchini et al., 2021; Filippi et al., 2000; Paci et al., 2024; Stedman et al., 1998).

Understanding cortical excitability modulation by cardiovascular activity is critical given the rising clinical interest in neurostimulation technologies (Jiao et al., 2024; Michael & Kaur, 2021). Substantial intra- and inter-subject TMS variability contributes to unpredictable therapeutic outcomes in TMS treatments (Sommer et al., 2002; Vallence et al., 2015; Ziemann & Siebner, 2015). Accounting for interoceptive and autonomic influences on brain states could improve neurostimulation reliability and predictability (Feng et al., 2025; Iseger et al., 2017; Michael & Kaur, 2021).

Two methodological limitations may contribute to previous inconsistencies. First, studies have not systematically controlled for spinal cord excitability, despite evidence that baroreceptor activation modulates spinal and brainstem reflexes (Dworkin et al., 1994; Rau, Brody, et al., 1993). Second, most studies used random TMS timing with post-hoc selection of MEPs within broad time intervals corresponding to ventricular systole and diastole (Al et al., 2023; Paci et al., 2024). Real-time ECG-TMS synchronization across several cardiac phases would enable precise temporal mapping of heart-driven neural excitability dynamics.

In the present study we analyzed spinal cord excitability changes during cardiac cycle using ECG-triggered trans-spinal magnetic stimulation. Real-time ECG monitoring and adaptation to heart rate changes enabled precise temporal targeting at five different moments within the systolic and diastolic phases with low timing variability. We compared the obtained pattern of spinal-MEP dynamics with the timings of different cardiovascular and cardiac-driven neural events in order to systematically analyze the potential mechanisms underlying the observed phenomenon. The study results may inform the optimization of TMS-based rehabilitation protocols for sensorimotor impairments following stroke, spinal cord injury and psychiatric disorders and should be considered in future investigations of cardiac-dependent corticospinal excitability.

## Methods

### Participants

Twenty healthy volunteers (10 females; mean age ± s.d. = 22.2 ± 4 years), all self-identified as right-handed, participated in the study. Participants were instructed to maintain 6–9 h of sleep prior to the experimental day, refrain from alcohol or medication for 24 h before the experiment, and avoid caffeine intake for at least 2 h prior to the procedure. Screening was performed to exclude contraindications to TMS in accordance with established safety guidelines (Rossi et al., 2009). The experimental protocol was approved by the Institutional Review Board of the Skolkovo Institute of Science and Technology (Approval No. 11, dated October 17, 2023). Each participant provided written informed consent prior to participating in the study.

### Experimental Design

Each participant completed a single experimental session lasting 45–60 min. Participants were seated in a chair designed to ensure stable and comfortable positioning throughout the experiment. The session consisted of three consecutive runs of transspinal magnetic stimulation (TSMS), each lasting 244 ± 43 s depending on the participant’s heart rate. Resting intervals were organized between runs.

Surface EMG and electrocardiography (ECG) were recorded simultaneously to assess spinal cord excitability across different phases of the cardiac cycle. EMG activity was recorded from the right hand muscles to capture TSMS-induced MEPs. Participants were instructed to remain fully relaxed and to avoid voluntary muscle activity to ensure that EMG signals reflected only TSMS-induced responses.

Surface bipolar electrodes were used to record activity from three muscles: *m*.*flexor digitorum superficialis* (FDS), *m*.*first dorsal interosseous* (FDI), and *m*.*abductor pollicis brevis* (APB). Active electrodes were placed over the muscle belly, reference electrodes over the tendon area. A ground electrode was placed on the left wrist.

ECG was recorded in standard Einthoven lead II configuration with electrodes placed on the right arm and left leg, providing reliable R-wave detection. Both EMG and ECG signals were digitized at 5 kHz using an NVX-52 amplifier (Medical Computer Systems, Russia). Electrode–skin impedance was maintained below 5 kΩ.

### ECG-triggered magnetic stimulation of spinal cord

Single biphasic magnetic pulses were delivered over the right side of the cervical spinal cord at the C6–T1 vertebral level using a figure-of-eight coil (d = 70 mm) connected to a Neuro-MSX stimulator (Neurosoft, Russia). For each participant, the optimal stimulation site (hotspot) was identified individually over the posterior midline at the C6-T1 vertebral level, where stimulation induced maximal FDS responses. The coil was positioned at 90° relative to the spinal axis to ensure targeted stimulation of segments related to right hand muscles (Fig. 1). With this orientation, the magnetic pulse was expected to activate dorsal roots (C6-C8 nerves) and interneurons in the posterior horns, thereby eliciting motor responses in the right hand via multisynaptic pathways (Carra et al., 2022; Fernandes et al., 2019). Stimulation intensity was set to 110% of the resting motor threshold for FDS, defined as the percentage of maximum stimulator output at which at least 5 out of 10 pulses evoked motor potentials exceeding 50 μV.

**Figure 1.**
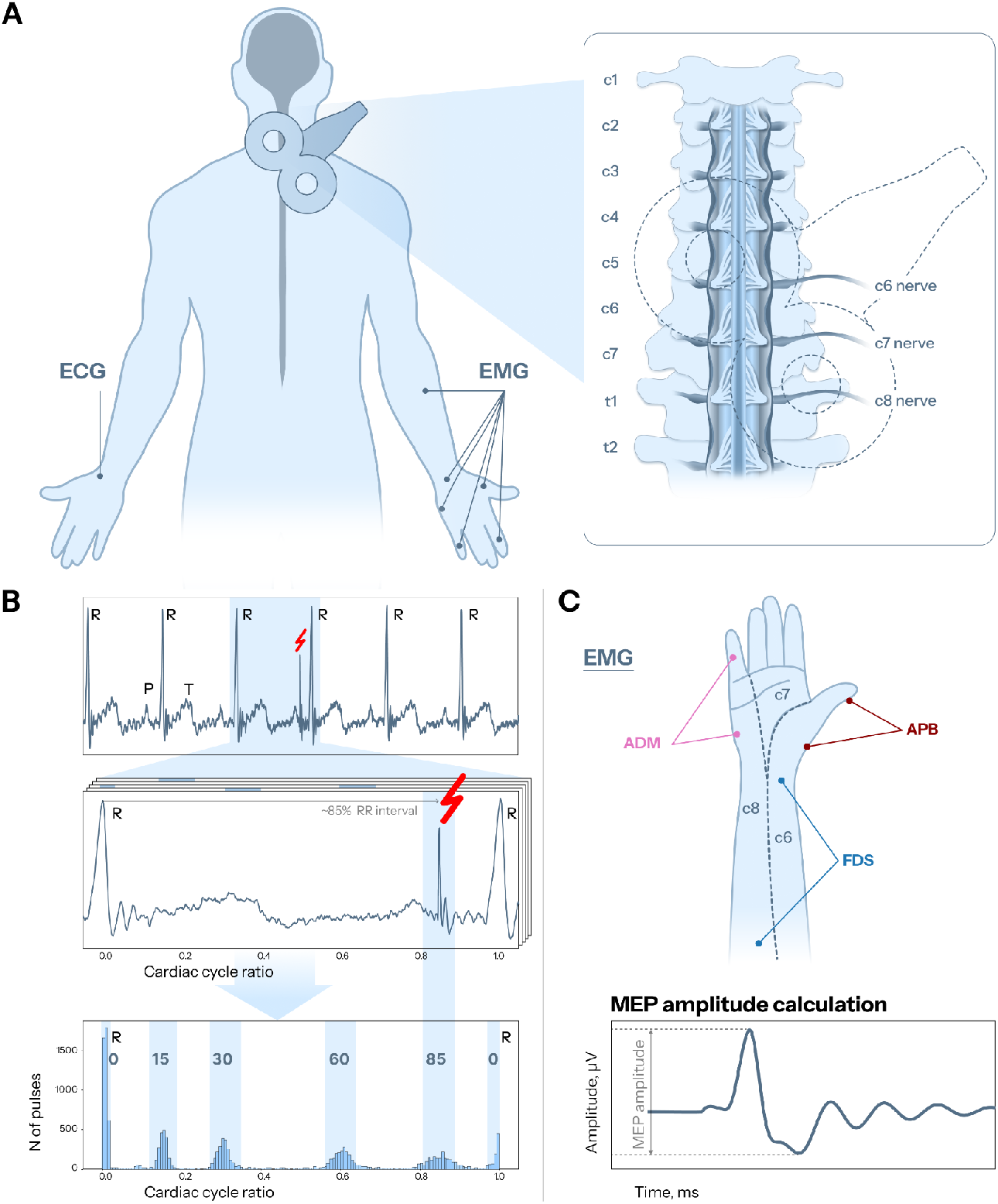
Methodology of ECG-triggered trans-spinal magnetic stimulation. (A) The magnetic stimulator figure-of-eight coil was positioned to activate C6-C8 dorsal root nerves and corresponding posterior horns, eliciting contractions specifically in fingers of the right hand. (B) ECG activity was recorded for real-time TSMS synchronization. The algorithm detected R-peaks and triggered stimulation at one of five preset time points corresponding to 0%, 15%, 30%, 60%, and 85% of the cardiac cycle. Raw ECG signal shows that at least two cardiac cycles passed between consecutive stimulations. The zoomed RR interval shows TSMS-induced artifacts in the ECG signal. In this example, stimulation was delivered at 85% of the cardiac cycle phase. Within a post-hoc analysis the exact relative phase of each stimulation was calculated. To minimize phase variability and create consistent trial groups, trials were stratified into bins defined as target phase ± 5% of the cardiac cycle. (C) EMG signals were recorded to obtain TSMS-evoked MEPs from the *m*.*flexor digitorum superficialis (FDS), m*.*first dorsal interosseous (FDI)*, and *m*.*abductor pollicis brevis (APB)* muscles. MEP amplitude was calculated as peak-to-peak amplitude.

During each run, magnetic pulses were delivered automatically using a custom Arduino-based device that recorded the ECG signal, detected R-peaks, and triggered the TMS machine via the synchronization port. Five R-peak–locked intervals corresponding to 0%, 15%, 30%, 60% and 85% of the cardiac cycle were preset. For a RR interval of 1000 ms, these time points corresponded to 0, 150, 300, 600 and 850 ms after the R peak. Prior to each stimulation, the delay was dynamically adjusted based on the duration of the preceding RR interval. The synchronization system enabled near real-time TSMS–ECG coupling, with delays between R-peak detection and stimulation onset not exceeding 60 μs for the 0% phase.

To avoid temporal regularity in stimulation sequence, consecutive pulses were separated by at least two cardiac cycles. The order of the stimulation phases was pseudorandomized to prevent 0% trials from following 85% trials (due to their close temporal proximity to avoid too frequent stimulation). 20–25 pulses per phase were collected per run, resulting in 60–75 total pulses per phase per participant.

## Data analysis

### Signal preprocessing

Data preprocessing and analysis were performed using Python packages MNE-Python (v1.6.0) (Gramfort et al., 2013), SciPy (v1.8.1) (Virtanen et al., 2020), Pingouin (v0.5.4) (Vallat, 2018) and Statsmodels (v0.14.1) (Seabold & Perktold, 2010). Raw signals were notch-filtered at 50 Hz and its harmonics up to 250 Hz to suppress powerline noise. EMG signal was band-pass filtered in 0.5 - 250 Hz range. All signals were epoched from −3 to +3 s relative to the TSMS pulse, which was marked online during recording.

For ECG, R peaks were identified using a peak detection algorithm (*find_peaks, SciPy*) with an adaptive minimum peak distance to account for inter-individual differences in heart rate and stimulation artifact occurrence. For each epoch, RR intervals were extracted. For subsequent analyses, we considered the interval preceding stimulation (RRpre), the interval of stimulation (RR), and the subsequent interval (RRpost). The relative phase when stimulation was applied within the cardiac cycle was computed as (R0–TSMS)/(R1–R0), where R0 and R1 denote consecutive R peaks, and TSMS is the time of spinal stimulation.

EMG epochs were baseline-corrected using the 100 ms preceding stimulation and cropped from −30 to +100 ms relative to the TSMS pulse. For each trial, MEP peak-to-peak amplitude was calculated within the 5–50 ms post-stimulation window. Only MEPs exceeding 50 μV were included in analyses. Thus, for each trial both MEP amplitudes and corresponding cardiac cycle phase were obtained. To mitigate variability in phase values and form trial groups, all trials were stratified into bins based on RR ratio values (see Fig.1B). Thus, after cleaning and stratification procedures on average 50 values per phase per participant left for final analysis.

### Statistical analysis

A two-way repeated measures ANOVA with factors **PHASE** and **MUSCLE** was used to assess effects of cardiac cycle phase on MEP amplitudes. Post-hoc analysis was performed using t-test for related samples. Significance level was set to p<0.05.

To evaluate the influence of preceding and subsequent RR intervals and to capture potential nonlinear relationships between excitability and cardiac phase, a linear mixed model (LMM) was fitted with peak-to-peak MEP amplitude (p2p) as the dependent variable. Fixed effects included the RR ratio (ratio), its quadratic term (ratio^2^), the pre-stimulation RR interval (RRpre), the stimulation RR interval (RR), and the post-stimulation RR interval (RRpost). A random intercept for the subject accounted for between-participant differences. The model was implemented in statsmodels (*mixedlm* function) as:

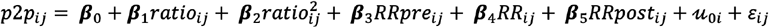

where u0i ∼ N(0, σu^2^) is the subject-level random intercept and εij ∼ N(0, σ^2^) is the residual error. Notably, for LMM fit we used all trials without binning stratification (see Fig.1B) because the model directly matched individual MEP amplitudes with exact cardiac phase values, enabling continuous analysis of phase-excitability relationships.

## Results

Motor evoked potential amplitudes were analyzed across all collected trials to assess cardiac cycle phase effects. Repeated-measures ANOVA results are shown in Table 1. A significant main effect of PHASE effect on MEP amplitudes was revealed (F = 3.9, p = 0.006). The main effect of MUSCLE did not reach statistical significance (F = 3.1, p = 0.054), though the observed trend may reflect higher MEP amplitudes in the FDS muscle compared to other recorded muscles (Fig.2A).

**Table 1.**
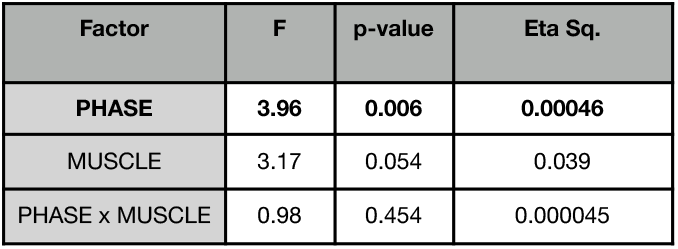
RM-ANOVA results for MEPs amplitude dynamics.

| Factor | F | p-value | Eta Sq. |
| --- | --- | --- | --- |
| PHASE | 3.96 | 0.006 | 0.00046 |
| MUSCLE | 3.17 | 0.054 | 0.039 |
| PHASE x MUSCLE | 0.98 | 0.454 | 0.000045 |

Figures 2B,C show MEP amplitude differences across cardiac phases. Maximal MEP amplitudes were obtained when TSMS was applied during diastolic intervals. Post-hoc analysis was focused on FDS MEPs, as this muscle was the target and exhibited higher MEP amplitudes, enabling reliable excitability estimation. T-tests revealed that MEPs obtained at 60% of the cardiac cycle were significantly higher than those during systolic phases (0% vs 60%: t = 2.6, p = 0.008; 15% vs 60%: t = 2.1, p = 0.025). Comparisons with 30% and 80% phases did not reach statistical significance (30% vs 60%: t = 1.45, p = 0.08; 80% vs 60%: t = 1.3, p = 0.10).

**Figure 2.**
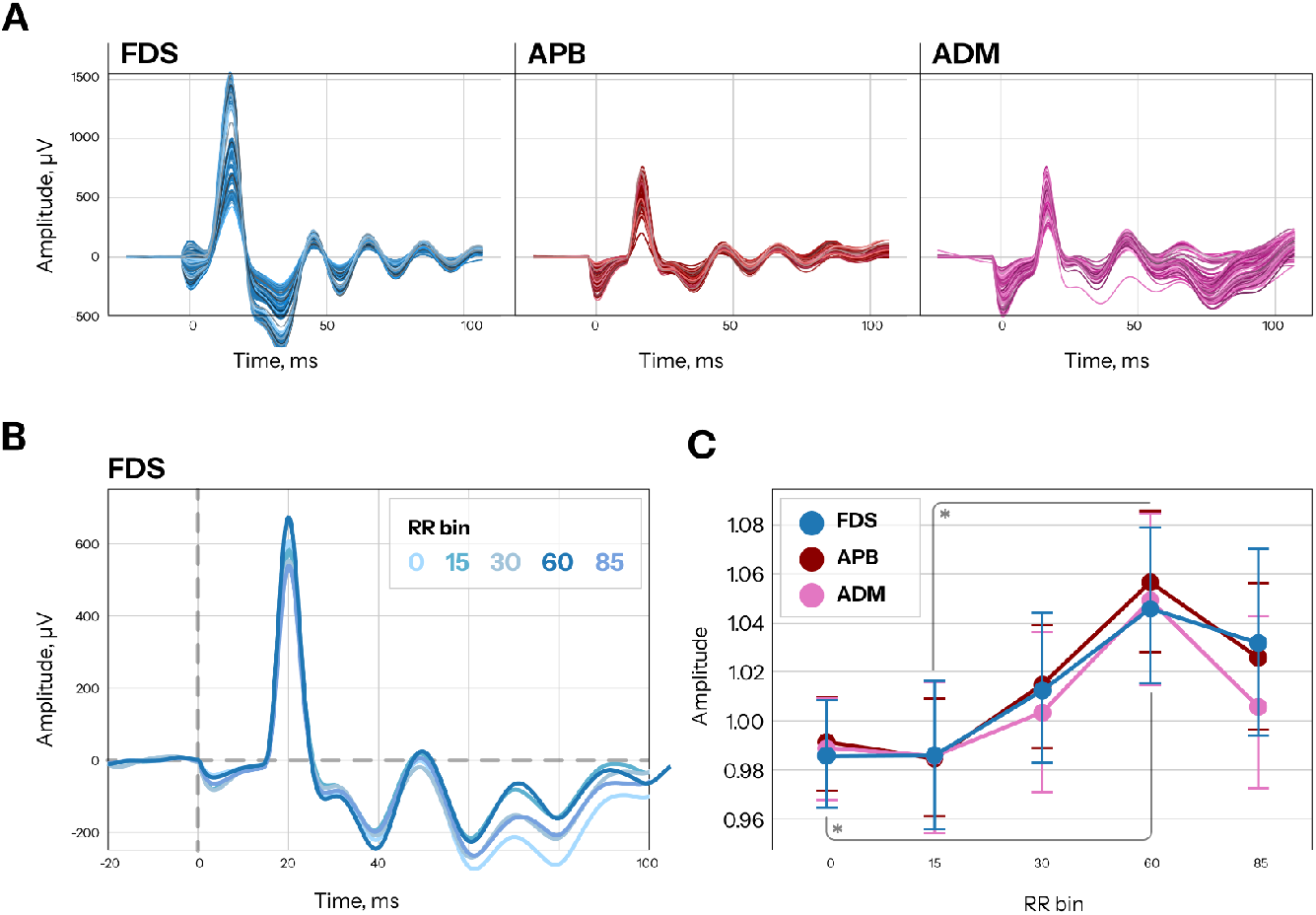
Cervical TSMS-induced MEPs and spinal cord excitability dynamics during the cardiac cycle. (A) Single-participant examples of MEPs recorded from different muscles. (B) Averaged MEPs from the FDS muscle grouped by cardiac cycle phase. (C) Group dynamics of MEP amplitudes for right-hand muscles across cardiac cycle phases. Mean and 95% CI are shown. Maximal responses to TSMS occurred at 60%. For visualization purposes, to account for between-subject and between-muscle variability, MEP amplitudes were normalized by dividing by the average amplitude for each subject/muscle combination.

To examine RR interval duration affects MEP amplitude dynamics, we analyzed single-trial data using a linear mixed model. Notably, all collected trials The model was fit to predict MEP amplitude based on cardiac cycle phase (and its quadratic term) and three RR interval durations: preceding (RRpre), current (RR), and following (RRpost) the excitability probe.

The model converged successfully under REML estimation. Analysis revealed a significant effect of RR phase on MEP amplitude (β = 67.93, SE = 32.00, z = 2.12, p = 0.034), while the quadratic phase effect did not reach significance (β = −59.40, SE = 34.98, z = −1.70, p = 0.09).

RRpre showed a significant negative association with MEP amplitude (β = −107.24, SE = 42.37, z = −2.53, p = 0.011), indicating that longer preceding RR intervals predicted reduced MEP amplitudes. Neither current RR (β = −62.23, SE = 49.99, z = −1.24, p = 0.213) nor RRpost (β = 21.58, SE = 32.54, z = 0.66, p = 0.507) reached significance level.

To illustrate the observed effects, all MEPs were stratified into groups based on preceding RR interval (RRpre) duration to separate responses following short versus long cardiac cycles. Based on RRpre distribution using a binning procedure we retained MEPs corresponding to the 5th-30th percentiles (short RRpre) and 70th-95th percentiles (long RRpre), excluding the extreme 5th and 95th percentiles as potentially noisy. The same binning procedure was performed based on RRpost distribution.

Figure 3 shows the binning results for the FDS muscle. MEP dynamics differed between stratified groups. The previously observed effect of diastolic MEPs increase is more pronounced in trials following short RRpre intervals (Fig.3A).

**Figure 3.**
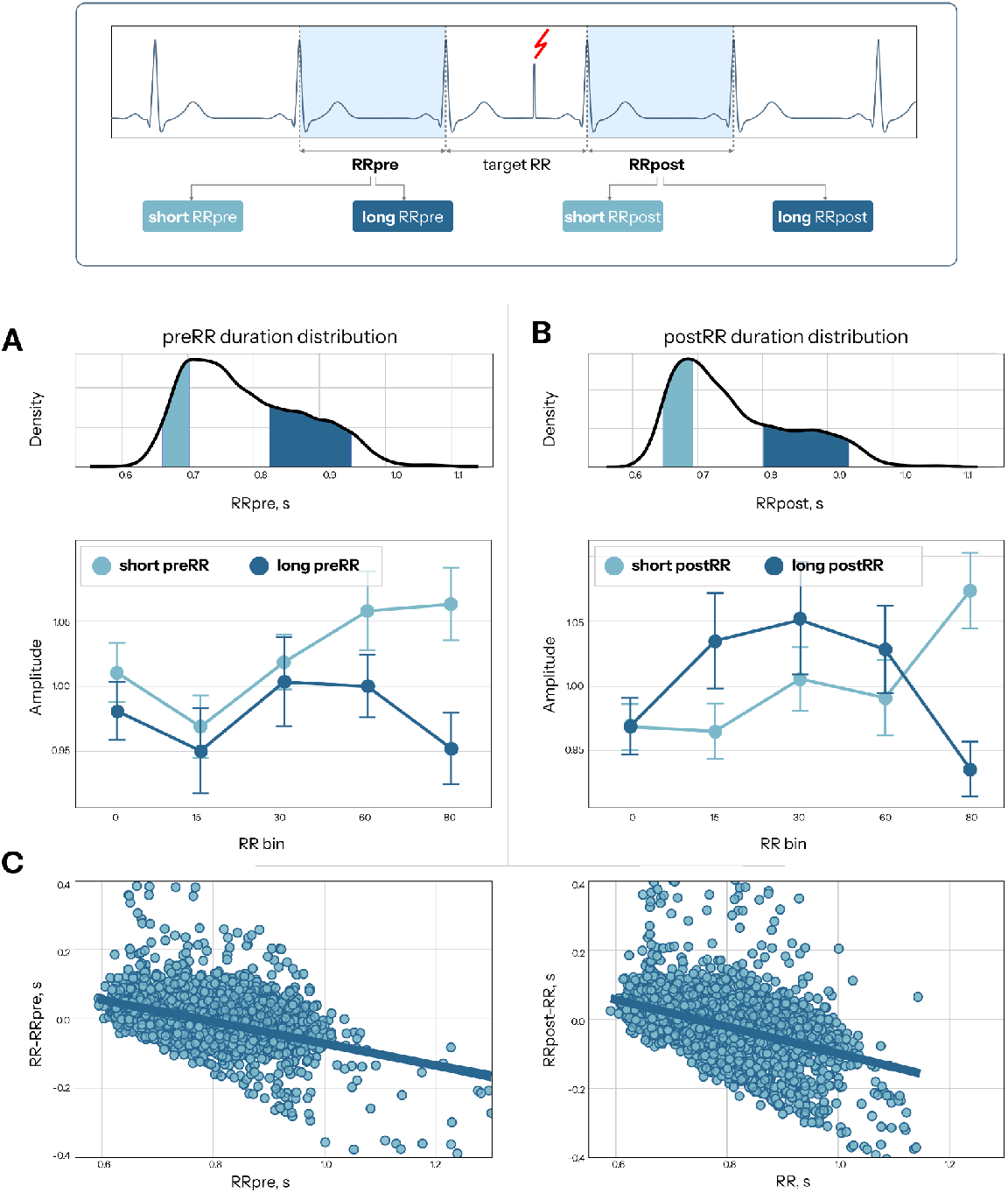
Spinal cord excitability dynamics during cardiac cycle in trials grouped based on proceeding (RRpre, A) or following (RRpost, B) RR interval duration. Mean and 95% CI are shown. RRpre and RRpost distributions are presented above corresponding graphs, highlighted regions indicate short and long interval subsamples. (C) Relationships between preceding RR duration and relative change in next RR. There is a tendency for decrease in RR after longer RR. RR intervals collected from all participants are shown.

Previous studies demonstrated that cortical stimulation affects heart rate in a cardiac cycle phase-dependent manner (Al et al., 2023). We analyzed cervical TSMS effects on RR intervals using two-way RM ANOVA with factors TIME (RRpre/RR/RRpost) and PHASE. Results revealed shortening in cardiac interval following the stimulation, with a significant main effect of TIME (F = 31.7, p < 0.001, η^2^ = 0.019). Neither the main effect of stimulation PHASE nor the TIME × PHASE interaction reached significance (Fig.4).

**Figure 4.**
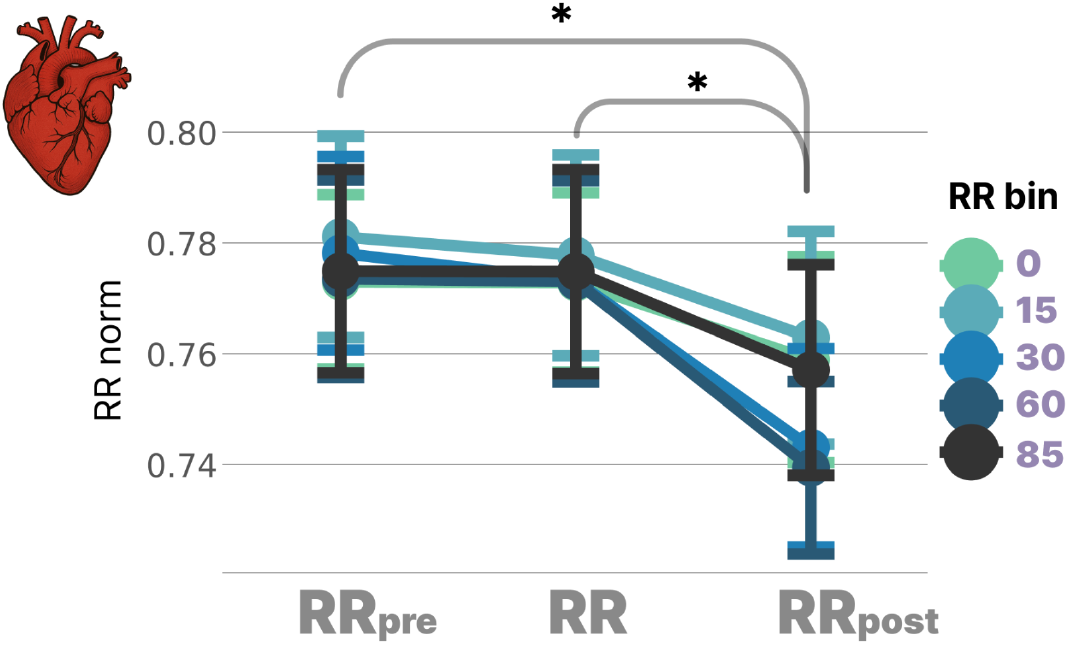
RR interval duration changes induced by cervical TSMS. Heart rate was accelerated after TSMS pulse, manifested as decreased RRpost duration. This effect was independent of a cardiac cycle phase in which stimulation was delivered. ^*^p < 0.0001, t-test.”

Since no stimulation phase effects were observed, pairwise comparisons used all trials averaged within subjects. Post-hoc t-tests revealed significant decreases in RRpost compared to both RR (t = 7.32, p < 0.0001) and RRpre (t = 5.43, p < 0.0001).

## Discussion

Our study explored spinal cord excitability modulation during the cardiac cycle with high temporal resolution using an adaptive ECG-triggered stimulation system. This approach enabled analysis of MEP dynamics at various points within the cardiac cycle, providing accurate excitability assessment beyond the traditional two-phase comparisons used in previous studies. Analysis of five time points covering systolic and diastolic events revealed that late diastole (60% of RR interval) was characterized by higher MEP amplitudes, indicating increased spinal excitability during this period.

### Central neural system activity depends on cardiac cycle

The interactions between the brain and the cardiovascular system has been the focus of many studies, with particular interest in how cardiac activity affects cortical excitability and brain functions. The brain is hypothesised to predict interoceptive signals during the systolic phase to suppress them from conscious perception while maintaining sensitivity to detect mismatches between predicted and actual bodily states (Ainley et al., 2016; Fló et al., 2024; Heck et al., 2016; Zaccaro et al., 2025). During the systolic phase increased “attention” to cardiac signal processing leads to attenuated somatosensory perception, resulting in reduced responses to external stimuli (Al et al., 2020; Motyka et al., 2019). At the same time, compensatory facilitation of motor functions during the systolic phase provides survival advantages through enhanced fight or flight readiness, suggesting distinct optimal windows for action versus perception throughout the cardiac cycle.

Our study, though, found higher spinal excitability during the diastole compared to systole, which contrasts with recent findings of increased M1 excitability during systole (Al et al., 2023). Since M1-evoked MEPs reflect combined cortical and spinal excitability, Al et al.’s results would predict either increased spinal MEPs during systole or no cardiac cycle effects.

At the same time our findings are consistent with previous reports of reduced spinal reflexes and pain perception during systole (Dworkin et al., 1994; Edwards et al., 2001; Rau, Brody, et al., 1993; Rau, Pauli, et al., 1993).

Thus, we propose that M1 and spinal excitability may fluctuate independently during the cardiac cycle. Spinal cord excitability appears modulated by baroreflex activity, while (Al et al., 2023) suggest M1 modulation is due to direct cardiac afferents that provide rapid systolic facilitation during the pre-ejection period, before pressure wave activates baroreceptors (Schulz et al., 2009).

Many TMS studies have failed to demonstrate the cardiac cycle modulation of M1 excitability, concluding that either cardiac activity does not influence cortical excitability or attributing null results to non-optimal stimulation timing. Our results suggest an alternative explanation: if motor cortex and spinal cord excitability fluctuate in antiphase during the cardiac cycle, M1 TMS induced MEPs may fail to detect cardiac cycle effects due to opposing modulations canceling each other out. Future studies should therefore focus on cortical excitability assessment via TEPs (Al et al., 2023).

### Mechanisms underlying cardiac cycle modulation of Spinal-MEPs

For illustrative purposes, all significant events occurring during the cardiac cycle were compiled from different articles and organized in Fig.5. Since in original articles some graphs were presented as percentages of the RR interval while others were reported in milliseconds after the R-peak, temporal adjustments were made based on the heart rate reported in each study or, when unreported, normalized to 60 BPM to standardize all data to the same temporal scale.

**Figure 5.**
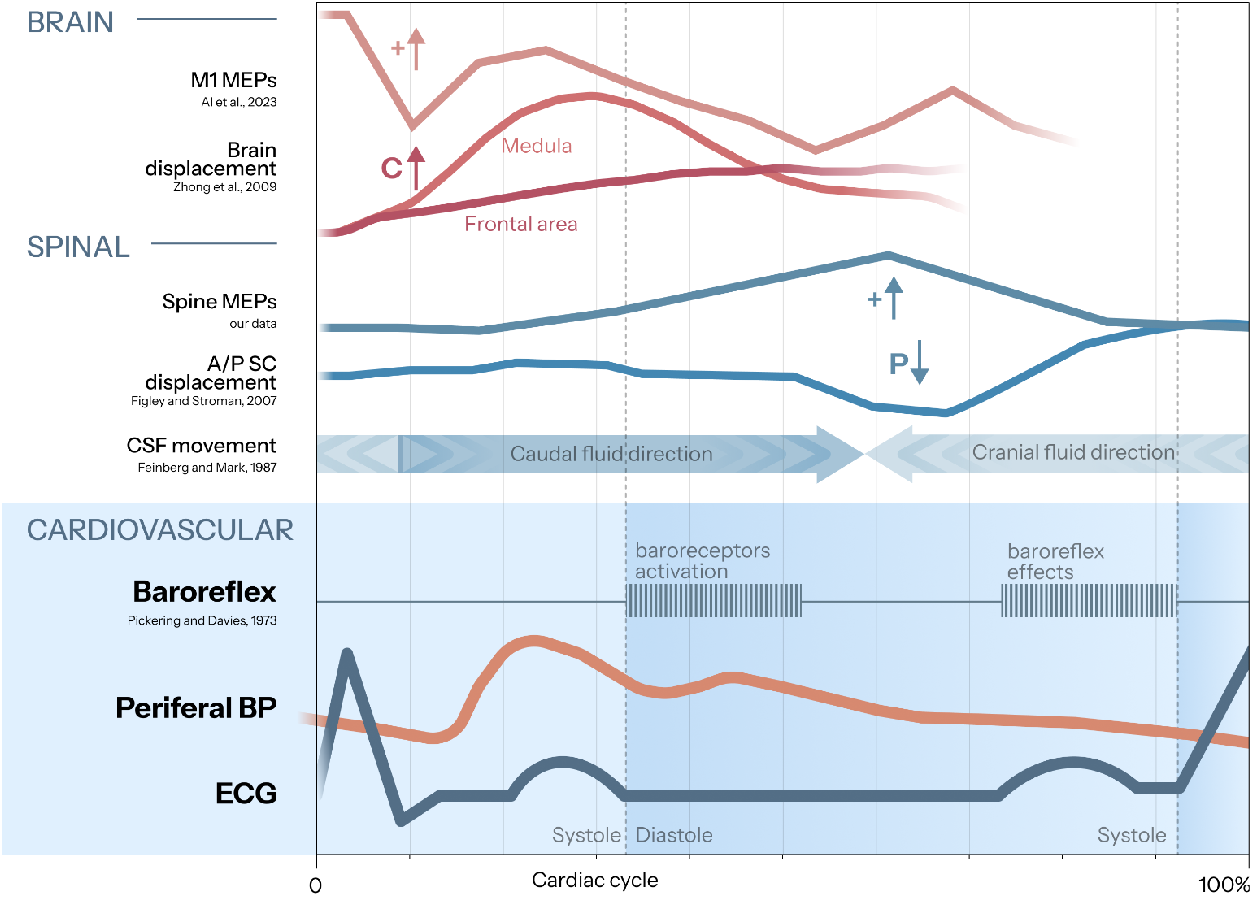
Temporal alignment of Cardiovascular and Ventral nervous system processes during the cardiac cycle. The ECG signal, peripheral blood pressure (BP), and baroreflex activity reflect cardiovascular system events (Borst & Karemaker, 1983; Pickering & Davies, 1973). Spinal cord events include cerebrospinal fluid (CSF) flow at the foramen magnum level (see (Feinberg & Mark, 1987), anterior-posterior (A/P) displacement of spinal cord (see (Figley & Stroman, 2007), and spinal-MEPs (our data). Corticospinal tract excitability is represented by primary motor cortex M1-MEPs dynamics (see (Al et al., 2023). Brain displacement graphs show movements of the medulla and frontal cortex toward the cranium (Zhong et al., 2009). Minor temporal adjustments were made to synchronize all graphs and demonstrate phase relationships across different processes. Arrow pictograms indicate axes directions: “+” positive, “P” posterior, “C” cranial. The horizontal axis represents one cardiac cycle (RR interval).

Two hypotheses may explain the current findings. First, the observed MEP dynamics may result from cardiac cycle-induced spinal cord displacement. The spinal cord undergoes rhythmic movement due to cerebrospinal fluid pulsatility (Feinberg & Mark, 1987). Any shifts in coil-to-tissue distance would result in induced current changes proportional to the inverse square of the distance (1/distance^2^). Detailed profile of spinal cord displacement throughout the cardiac cycle was documented in (Figley et al., 2008; Figley & Stroman, 2007).

Given the stimulation coil was positioned directly on the participant’s spine, anterior-posterior spinal cord movement may affect stimulation efficacy: posterior displacement brings neural tissue closer to the coil, increasing magnetic field intensity, while anterior displacement reduces field strength. The cervico-thoracic junction (C4-T1) exhibits maximal cardiac-related anterior-posterior displacement, with amplitudes exceeding 1 mm during normal breathing (Winklhofer et al., 2014). Comparison of anterior-posterior movement profiles at C4-C6 levels reported by (Figley & Stroman, 2007) with our MEP amplitude dynamics revealed notable correspondence between MEP increases and maximal spinal cord proximity to the dorsal surface. (Fig.5).

Mechanical displacement of brain tissue may similarly explain increased M1 excitability in TMS studies. Although Al et al. (2023) rejected the mechanical displacement hypothesis after comparing M1-MEP dynamics with brain movement data, they used subcortical displacement (inverted U-shaped trajectory, see Fig. 5) rather than the cortical movement trajectory, which shows distinct pattern (Feinberg & Mark, 1987; Zhong et al., 2009). Cortical tissue moves toward the skull approximately 100 ms after the R-wave and returns caudally at 200 ms during systole (Feinberg & Mark, 1987), which could partially explain the systolic M1-MEP increases reported by (Al et al., 2023).

However, brain movement cannot explain other cardiac-related phenomena such as sensory attenuation, increased muscle force during systole, or recently described increased GABA-A interneurons activity in the motor cortex (Paci et al., 2024). Furthermore, if M1-MEP dynamics resulted solely from brain movement, other TMS studies would likely have detected similar cardiac cycle effects, yet (Al et al., 2023) are among the few to report such findings. These considerations suggest that other mechanisms beyond tissue displacement are required to fully explain the cardiac cycle effects on neural excitability.

Another physiological mechanism that might explain our findings is baroreflex activity, which effects span beyond cardiovascular regulations (Duschek et al., 2013). During systole, increased blood pressure activates carotid sinus baroreceptors approximately 300 ms after the R-peak, with an additional ∼400 ms required for reflex loop completion and cardiac modulation (∼600-800 ms from R-peak to baroreflex manifestation (Borst & Karemaker, 1983; Pickering & Davies, 1973).

Additionally, baroreflex has been shown to have inhibitory effects on central neural circuits at both cerebral and spinal levels (Dworkin et al., 1994; Rau, Pauli, et al., 1993). Baroreceptor stimulation increases slow EEG activity (Braun et al., 2025; Cole, 1989) and reduces spinal reflexes (Rau, Brody, et al., 1993).

The inhibitory effects on the nervous system appear faster than cardiac effects and manifests during the first half of the cardiac cycle (Al et al., 2023; Pickering & Davies, 1973). Conversely, during diastole when baroreceptor activity decreases, spinal cord reactivity increases (Rau, Brody, et al., 1993). This pattern aligns with our findings of increased spinal MEPs during late diastole, before atrial systole.

To further test the baroreflex hypothesis, we analyzed the effects of RR interval duration on MEPs across three intervals: preceding (RRpre), current (RR), and subsequent (RRpost) to TSMS application. We hypothesized that shorter preceding intervals would produce higher baroreflex activation, leading to lower baroreflex activity in the subsequent cycle and thus higher spinal excitability. Conversely, longer preceding intervals were expected to yield stronger baroreflex activation during the TSMS cycle, resulting in reduced spinal excitability. For illustration of the relationship between two consecutive RR intervals, see Fig.4C.

The obtained results supported this hypothesis: short RRpre intervals were associated with stronger diastolic MEP increases. LMM analysis confirmed that preceding interval duration significantly affected spinal excitability, supporting baroreflex modulation of spinal circuits. However, current RR and RRpost showed no significant effects. Future experiments with arterial pressure recordings and controlled carotid sinus stimulation using neck suction techniques (Rau, Brody, et al., 1993) would directly validate these baroreflex-mediated effects.

### Spinal cord stimulation affects heart rate

Future experiments should also account for neurostimulation effects on cardiac activity. We found that spinal stimulation decreased subsequent RR interval duration, independent of the cardiac phase during which TSMS was applied. This contrasts with previous TMS findings where Al et al. reported longer RR intervals when stimulation was applied during the systolic phase, with effects observed on the current rather than subsequent interval (Al et al., 2023).

Several mechanisms could explain our findings. TSMS magnetic field could directly stimulate carotid sinus baroreceptors or baroreflex afferent pathways, triggering heart rate changes in the subsequent cardiac cycle. Alternatively, the auditory startle response from TSMS acoustic noise could accelerate heart rate, though this mechanism would be expected to affect the current RR interval, particularly for stimuli applied at the R-peak (0% phase).

The ability of single-pulse spinal stimulation to modulate cardiac activity has potential clinical applications. If TSMS can effectively modulate baroreflex function, this approach might benefit treatment of cardiovascular (Howard-Quijano et al., 2017; Kuwabara et al., 2023) and psychiatric disorders characterized by reduced baroreflex sensitivity, including depression, anxiety, schizophrenia, and fatigue syndromes (Duschek et al., 2013; Kaur et al., 2020; Michael & Kaur, 2021; Peckerman et al., 2003; Reyes del Paso et al., 2021).

## Conclusion

This study demonstrates that spinal cord excitability fluctuates during the cardiac cycle in patterns distinct from previously reported cortical dynamics. This mismatch may explain why M1-TMS studies typically fail to detect cardiac cycle effects on MEPs, as opposing cortical and spinal modulations could cancel each other out. Our analysis revealed peak spinal excitability during diastole, coinciding with reduced baroreflex activity, with dynamics further modulated by preceding RR interval duration—supporting baroreflex involvement in these effects. However, future studies are still needed to define the relative contributions of neural tissue pulsation versus neurophysiological mechanisms to cardiac cycle effects on neural excitability.

### Limitations and Future Directions

Several methodological issues limit the interpretation of our findings. First, our R-peak-based triggering system still introduced temporal variability due to beat-to-beat heart rate fluctuations. The system calculated stimulation delays from the R-peak, and this created a variability between intended and actual cardiac phase stimulation was applied: early phases (15-30%) maintained precision, while late phases (60-85%) exhibited greater deviation from target timings (see Fig.1). Although offline MEP selection based on the actual cardiac phase minimized this drawback, it reduced the number of analyzable trials.

Second, we did not control for respiratory influences on cardiovascular dynamics and spinal cord displacement. Respiratory modulation of blood pressure, neural excitability, and mechanical tissue motion requires systematic investigation in future studies (Engelen et al., 2024; Heck et al., 2016; Winklhofer et al., 2014).

Third, our findings are limited to cervical spinal segments. Given that caudal spinal regions exhibit distinct pulsatility profiles with reduced displacement amplitudes (Figley et al., 2008; Figley & Stroman, 2007), future studies should examine multiple spinal levels to determine the generalizability of cardiac cycle effects on spinal excitability.

## Author Contributions

NS contributed to the conception and design of the experiments; data collection and analysis; results interpretation; drafting of the manuscript; editing and revision of the manuscript. AP, EM, RT, and PM equally contributed to the design of the experiments and data collection. AM: data analysis and results presentation. AB and LY contributed to the editing and revision of the manuscript. MK: design of the experiments, AK supervised the study.

## Data Availability

The data collected and analyzed during the current study is available from the corresponding author on reasonable request.

## Acknowledgments

This work was supported by the scientific project of the state assignment of Lomonosov Moscow State University, No. 121032300070-1.

## Disclosures

The authors declare no competing interests.

## Notes

### Competing Interest Statement

The authors have declared no competing interest.

## References

Ainley, V., Apps, M. A. J., Fotopoulou, A., & Tsakiris, M. (2016). ‘Bodily precision’: A predictive coding account of individual differences in interoceptive accuracy. Philosophical Transactions of the Royal Society B: Biological Sciences, 371(1708), 20160003. 10.1098/rstb.2016.0003

Al, E., Iliopoulos, F., Forschack, N., Nierhaus, T., Grund, M., Motyka, P., Gaebler, M., Nikulin, V. V., & Villringer, A. (2020). Heart–brain interactions shape somatosensory perception and evoked potentials. Proceedings of the National Academy of Sciences, 117(19), 10575–10584. 10.1073/pnas.1915629117

Al, E., Stephani, T., Engelhardt, M., Haegens, S., Villringer, A., & Nikulin, V. V. (2023). Cardiac activity impacts cortical motor excitability. PLOS Biology, 21(11), e3002393. 10.1371/journal.pbio.3002393

Bianchini, E., Mancuso, M., Zampogna, A., Guerra, A., & Suppa, A. (2021). Cardiac cycle does not affect motor evoked potential variability: A real-time EKG-EMG study. Brain Stimulation, 14(1), 170–172. 10.1016/j.brs.2020.12.009

Borst, C., & Karemaker, J. M. (1983). Time delays in the human baroreceptor reflex. Journal of the Autonomic Nervous System, 9(2–3), 399–409. 10.1016/0165-1838(83)90004-8

Braun, J., Patel, M., Woods, W., Keatch, C., Kameneva, T., & Lambert, E. (2025). Frontal and temporo-parietal changes in delta and alpha power accompany stress-induced vasoconstriction and blood pressure response. Journal of Neurophysiology, 133(6), 1815–1827. 10.1152/jn.00618.2024

Carra, R. B., Silva, G. D., Paraguay, I. B. B., Diniz De Lima, F., Menezes, J. R., Pineda, A. M., Nunes, G. A., Simões, J. D. S., França, M. C., & Cury, R. G. (2022). Controversies and Clinical Applications of Non-Invasive Transspinal Magnetic Stimulation: A Critical Review and Exploratory Trial in Hereditary Spastic Paraplegia. Journal of Clinical Medicine, 11(16), 4748. 10.3390/jcm11164748

Cole, R. J. (1989). Postural baroreflex stimuli may affect EEG arousal and sleep in humans. Journal of Applied Physiology, 67(6), 2369–2375. 10.1152/jappl.1989.67.6.2369

De Falco, E., Solcà, M., Bernasconi, F., Babo-Rebelo, M., Young, N., Sammartino, F., Tallon-Baudry, C., Navarro, V., Rezai, A. R., Krishna, V., & Blanke, O. (2024). Single neurons in the thalamus and subthalamic nucleus process cardiac and respiratory signals in humans. Proceedings of the National Academy of Sciences, 121(11), e2316365121. 10.1073/pnas.2316365121

Duschek, S., Werner, N. S., & Reyes Del Paso, G. A. (2013). The behavioral impact of baroreflex function: A review. Psychophysiology, 50(12), 1183–1193. 10.1111/psyp.12136

Dworkin, B. R., Elbert, T., Rau, H., Birbaumer, N., Pauli, P., Droste, C., & Brunia, C. H. (1994). Central effects of baroreceptor activation in humans: Attenuation of skeletal reflexes and pain perception. Proceedings of the National Academy of Sciences, 91(14), 6329–6333. 10.1073/pnas.91.14.6329

Edwards, L., Ring, C., McIntyre, D., & Carroll, D. (2001). Modulation of the human nociceptive flexion reflex across the cardiac cycle. Psychophysiology, 38(4), 712–718. 10.1111/1469-8986.3840712

Engelen, T., Schuhmann, T., Sack, A. T., & Tallon-Baudry, C. (2024). The cardiac, respiratory, and gastric rhythms independently modulate motor corticospinal excitability in humans. Neuroscience. 10.1101/2024.09.10.612221

Feinberg, D. A., & Mark, A. S. (1987). Human brain motion and cerebrospinal fluid circulation demonstrated with MR velocity imaging. Radiology, 163(3), 793–799. 10.1148/radiology.163.3.3575734

Feng, Z.-J., Martin, S., Numssen, O., Weise, K., Jing, Y., Gerardos, G., Martin, C., Hartwigsen, G., & Knösche, T. R. (2025). Target-Specificity and Repeatability in Neuro-Cardiac-Guided TMS for Heart-Brain Coupling (p. 2025.02.19.638988). bioRxiv. 10.1101/2025.02.19.638988

Fernandes, S. R., Salvador, R., De Carvalho, M., & Miranda, P. C. (2019). Electric Field Distribution during Non-Invasive Electric and Magnetic Stimulation of the Cervical Spinal Cord. 2019 41st Annual International Conference of the IEEE Engineering in Medicine and Biology Society (EMBC), 5898–5901. 10.1109/EMBC.2019.8857129

Figley, C. R., & Stroman, P. W. (2007). Investigation of human cervical and upper thoracic spinal cord motion: Implications for imaging spinal cord structure and function. Magnetic Resonance in Medicine, 58(1), 185–189. 10.1002/mrm.21260

Figley, C. R., Yau, D., & Stroman, P. W. (2008). Attenuation of Lower-Thoracic, Lumbar, and Sacral Spinal Cord Motion: Implications for Imaging Human Spinal Cord Structure and Function. American Journal of Neuroradiology, 29(8), 1450–1454. 10.3174/ajnr.A1154

Filippi, M. M., Oliveri, M., Vernieri, F., Pasqualetti, P., & Rossini, P. M. (2000). Are autonomic signals influencing cortico-spinal motor excitability?: A study with transcranial magnetic stimulation. Brain Research, 881(2), 159–164. 10.1016/S0006-8993(00)02837-7

Fló, E., Belloli, L., Cabana, Á., Ruyant-Belabbas, A., Jodaitis, L., Valente, M., Rohaut, B., Naccache, L., Rosanova, M., Comanducci, A., Andrillon, T., & Sitt, J. (2024). Predicting attentional focus: Heartbeat-evoked responses and brain dynamics during interoceptive and exteroceptive processing. PNAS Nexus, 3(12), pgae531. 10.1093/pnasnexus/pgae531

Galvez-Pol, A., Virdee, P., Villacampa, J., & Kilner, J. (2022). Active tactile discrimination is coupled with and modulated by the cardiac cycle. eLife, 11, e78126. 10.7554/eLife.78126

Gramfort, A., Luessi, M., Larson, E., Engemann, D. A., Strohmeier, D., Brodbeck, C., Goj, R., Jas, M., Brooks, T., Parkkonen, L., & Hämäläinen, M. (2013). MEG and EEG data analysis with MNE-Python. Frontiers in Neuroscience, 7. 10.3389/fnins.2013.00267

Heck, D. H., McAfee, S. S., Liu, Y., Babajani-Feremi, A., Rezaie, R., Freeman, W. J., Wheless, J. W., Papanicolaou, A. C., Ruszinkó, M., & Kozma, R. (2016). Cortical rhythms are modulated by respiration. Neuroscience. 10.1101/049007

Howard-Quijano, K., Takamiya, T., Dale, E. A., Kipke, J., Kubo, Y., Grogan, T., Afyouni, A., Shivkumar, K., & Mahajan, A. (2017). Spinal cord stimulation reduces ventricular arrhythmias during acute ischemia by attenuation of regional myocardial excitability. American Journal of Physiology-Heart and Circulatory Physiology, 313(2), H421–H431. 10.1152/ajpheart.00129.2017

Iseger, T. A., Padberg, F., Kenemans, J. L., Gevirtz, R., & Arns, M. (2017). Neuro-Cardiac-Guided TMS (NCG-TMS): Probing DLPFC-sgACC-vagus nerve connectivity using heart rate – First results. Brain Stimulation, 10(5), 1006–1008. 10.1016/j.brs.2017.05.002

Jammal Salameh, L., Bitzenhofer, S. H., Hanganu-Opatz, I. L., Dutschmann, M., & Egger, V. (2024). Blood pressure pulsations modulate central neuronal activity via mechanosensitive ion channels. Science, 383(6682), eadk8511. 10.1126/science.adk8511

Jiao, Y., Cheng, C., Jia, M., Chu, Z., Song, X., Zhang, M., Xu, H., Zeng, X., Sun, J.-B., Qin, W., & Yang, X.-J. (2024). Neuro-cardiac-guided transcranial magnetic stimulation: Unveiling the modulatory effects of low-frequency and high-frequency stimulation on heart rate. Psychophysiology, 61(10), e14631. 10.1111/psyp.14631

Kaur, M., Michael, J. A., Hoy, K. E., Fitzgibbon, B. M., Ross, M. S., Iseger, T. A., Arns, M., Hudaib, A.-R., & Fitzgerald, P. B. (2020). Investigating high- and low-frequency neuro-cardiac-guided TMS for probing the frontal vagal pathway. Brain Stimulation, 13(3), 931–938. 10.1016/j.brs.2020.03.002

Khoshnoud, S., Leitritz, D., Bozdağ, M. Ç., Igarzábal, F. A., Noreika, V., & Wittmann, M. (2024). When the Heart Meets the Mind: Exploring the Brain–Heart Interaction during Time Perception. Journal of Neuroscience, 44(34). 10.1523/JNEUROSCI.2039-23.2024

Konttinen, N., Mets, T., Lyytinen, H., & Paananen, M. (2003). Timing of Triggering in Relation to the Cardiac Cycle in Nonelite Rifle Shooters. Research Quarterly for Exercise and Sport, 74(4), 395–400. 10.1080/02701367.2003.10609110

Kuwabara, Y., Howard-Quijano, K., Salavatian, S., Yamaguchi, T., Saba, S., & Mahajan, A. (2023). Thoracic dorsal root ganglion stimulation reduces acute myocardial ischemia induced ventricular arrhythmias. Frontiers in Neuroscience, 17. 10.3389/fnins.2023.1091230

Larra, M. F., Finke, J. B., Wascher, E., & Schächinger, H. (2020). Disentangling sensorimotor and cognitive cardioafferent effects: A cardiac-cycle-time study on spatial stimulus-response compatibility. Scientific Reports, 10(1), 4059. 10.1038/s41598-020-61068-1

Michael, J. A., & Kaur, M. (2021). The Heart-Brain Connection in Depression: Can it inform a personalised approach for repetitive transcranial magnetic stimulation (rTMS) treatment? Neuroscience & Biobehavioral Reviews, 127, 136–143. 10.1016/j.neubiorev.2021.04.016

Motyka, P., Grund, M., Forschack, N., Al, E., Villringer, A., & Gaebler, M. (2019). Interactions between cardiac activity and conscious somatosensory perception. Psychophysiology, 56(10), e13424. 10.1111/psyp.13424

Mussini, E., Zaccaro, A., Perrucci, M. G., Costantini, M., & Ferri, F. (2025). Heartbeat on hold: Cortical processing of cardiac signals during motor preparation. NeuroImage, 316, 121299. 10.1016/j.neuroimage.2025.121299

Paci, M., Cardellicchio, P., Di Luzio, P., Perrucci, M. G., Ferri, F., & Costantini, M. (2024). When the heart inhibits the brain: Cardiac phases modulate short-interval intracortical inhibition. iScience, 27(3), 109140. 10.1016/j.isci.2024.109140

Palser, E. R., Glass, J., Fotopoulou, A., & Kilner, J. M. (2021). Relationship between cardiac cycle and the timing of actions during action execution and observation. Cognition, 217, 104907. 10.1016/j.cognition.2021.104907

Peckerman, A., LaManca, J. J., Qureishi, B., Dahl, K. A., Golfetti, R., Yamamoto, Y., & Natelson, B. H. (2003). Baroreceptor Reflex and Integrative Stress Responses in Chronic Fatigue Syndrome. Biopsychosocial Science and Medicine, 65(5), 889. 10.1097/01.PSY.0000079408.62277.3D

Pickering, T. G., & Davies, J. (1973). Estimation of the conduction time of the baroreceptor-cardiac reflex in man. Cardiovascular Research, 7(2), 213–219. 10.1093/cvr/7.2.213

Rau, H., Brody, S., Brunia, C. H. M., Damen, E. P. J., & Elbert, T. (1993). Activation of carotid baroreceptors inhibits spinal reflexes in man. Electroencephalography and Clinical Neurophysiology/Evoked Potentials Section, 89(5), 328–334. 10.1016/0168-5597(93)90072-W

Rau, H., Pauli, P., Brody, S., Elbert, T., & Birbaumer, N. (1993). Baroreceptor stimulation alters cortical activity. Psychophysiology, 30(3), 322–325. 10.1111/j.1469-8986.1993.tb03359.x

Reyes del Paso, G. A., Contreras-Merino, A. M., de la Coba, P., & Duschek, S. (2021). The cardiac, vasomotor, and myocardial branches of the baroreflex in fibromyalgia: Associations with pain, affective impairments, sleep problems, and fatigue. Psychophysiology, 58(5), e13800. 10.1111/psyp.13800

Rossi, S., Hallett, M., Rossini, P. M., & Pascual-Leone, A. (2009). Safety, ethical considerations, and application guidelines for the use of transcranial magnetic stimulation in clinical practice and research. Clinical Neurophysiology, 120(12), 2008–2039. 10.1016/j.clinph.2009.08.016

Schulz, A., Lass-Hennemann, J., Nees, F., Blumenthal, T. D., Berger, W., & Schachinger, H. (2009). Cardiac modulation of startle eye blink. Psychophysiology, 46(2), 234–240. 10.1111/j.1469-8986.2008.00768.x

Seabold, S., & Perktold, J. (2010). Statsmodels: Econometric and Statistical Modeling with Python. 92–96. 10.25080/Majora-92bf1922-011

Sommer, M., Wu, T., Tergau, F., & Paulus, W. (2002). Intra- and interindividual variability of motor responses to repetitive transcranial magnetic stimulation. Clinical Neurophysiology, 113(2), 265–269. 10.1016/S1388-2457(01)00726-X

Stedman, A., Davey, N. J., & Ellaway, P. H. (1998). Facilitation of human first dorsal interosseous muscle responses to transcranial magnetic stimulation during voluntary contraction of the contralateral homonymous muscle. Muscle & Nerve, 21(8), 1033–1039. 10.1002/(SICI)1097-4598(199808)21:8%253C1033::AID-MUS7%253E3.0.CO;2-9

Vallat, R. (2018). Pingouin: Statistics in Python. Journal of Open Source Software, 3(31), 1026. 10.21105/joss.01026

Vallence, A.-M., Goldsworthy, M. R., Hodyl, N. A., Semmler, J. G., Pitcher, J. B., & Ridding, M. C. (2015). Inter- and intra-subject variability of motor cortex plasticity following continuous theta-burst stimulation. Neuroscience, 304, 266–278. 10.1016/j.neuroscience.2015.07.043

Virtanen, P., Gommers, R., Oliphant, T. E., Haberland, M., Reddy, T., Cournapeau, D., Burovski, E., Peterson, P., Weckesser, W., Bright, J., Van Der Walt, S. J., Brett, M., Wilson, J., Millman, K. J., Mayorov, N., Nelson, A. R. J., Jones, E., Kern, R., Larson, E., … Vázquez-Baeza, Y. (2020). SciPy 1.0: Fundamental algorithms for scientific computing in Python. Nature Methods, 17(3), 261–272. 10.1038/s41592-019-0686-2

Winklhofer, S., Schoth, F., Stolzmann, P., Krings, T., Mull, M., Wiesmann, M., & Stracke, C. (2014). Spinal Cord Motion: Influence of Respiration and Cardiac Cycle. RöFo - Fortschritte Auf Dem Gebiet Der Röntgenstrahlen Und Der Bildgebenden Verfahren, 186(11), 1016–1021. 10.1055/s-0034-1366429

Zaccaro, A., Penna, F. della, Bubbico, F., Bayram, B., Parrotta, E., Perrucci, M. G., Costantini, M., & Ferri, F. (2025). Cardio-respiratory interactions in interoceptive perception: The role of heartbeat-modulated cortical oscillations (p. 2025.04.14.648690). bioRxiv. 10.1101/2025.04.14.648690

Zhong, X., Meyer, C. H., Schlesinger, D. J., Sheehan, J. P., Epstein, F. H., Larner, J. M., Benedict, S. H., Read, P. W., Sheng, K., & Cai, J. (2009). Tracking brain motion during the cardiac cycle using spiral cine-DENSE MRI: Track pulsatile brain motion using DENSE MRI. Medical Physics, 36(8), 3413–3419. 10.1118/1.3157109

Ziemann, U., & Siebner, H. R. (2015). Inter-subject and Inter-session Variability of Plasticity Induction by Non-invasive Brain Stimulation: Boon or Bane? Brain Stimulation, 8(3), 662–663. 10.1016/j.brs.2015.01.409

